# dream: Powerful differential expression analysis for repeated measures designs

**DOI:** 10.1101/432567

**Authors:** Gabriel E. Hoffman, Panos Roussos

**Affiliations:** Pamela Sklar Division of Psychiatric Genomics, Icahn School of Medicine at Mount Sinai, New York, NY, USA; Icahn Institute for Data Science and Genomic Technology, Icahn School of Medicine at Mount Sinai, New York, NY, USA; Department of Genetics and Genomic Sciences, Icahn School of Medicine at Mount Sinai, New York, NY, USA; Department of Psychiatry, Icahn School of Medicine at Mount Sinai, New York, NY, USA; Mental Illness Research, Education, and Clinical Center (VISN 2 South), James J. Peters VA Medical Center, Bronx, NY, USA

## Abstract

Large-scale transcriptome studies with multiple samples per individual are widely used to study disease biology. Yet current methods for differential expression are inadequate for cross-individual testing for these repeated measures designs. Most problematic, we observe across multiple datasets that current methods can give reproducible false positive findings that are driven by genetic regulation of gene expression, yet are unrelated to the trait of interest. Here we introduce a statistical software package, dream, that increases power, controls the false positive rate, enables multiple types of hypothesis tests, and integrates with standard workflows. In 12 analyses in 6 independent datasets, dream yields biological insight not found with existing software while addressing the issue of reproducible false positive findings. Dream is available within the variancePartition Bioconductor package (http://bioconductor.org/packages/variancePartition).

## INTRODUCTION

Transcriptome profiling and comparison of gene expression levels are a widely used genomic technique in biomedical research. In a typical study, a researcher collects gene expression by RNA-seq or microarray from multiple samples and performs differential expression analysis between subsets of samples that differ in cell/tissue type, environmental conditions, stimuli, genotype, or disease state. A range of statistical methods have been developed for this purpose and give state-of-the-art performance on this typical study design (1, 2, 3, 4, 5, 6).

Recent advances in the scale of transcriptomic and, more generally, functional genomic studies have enabled assaying individuals from multiple tissues (7, 8), brain regions (9, 10), cell types (11), time points (12, 13, 14) or induced pluripotent stem cell (iPSC) lines (15, 16, 17, 18, 19, 20). These studies with multiple samples from each individual can test region- or context-specific effects, but can also increase the statistical power to detect effects that are shared by multiple replicates (15, 21, 22). Collecting replicates also enables samples from the same individual to be processed in multiple technical batches in order to decouple biological from technical variation in gene expression (23). Finally, collecting replicates can also be beneficial when gene expression measurements are noisy, when expression is dynamic or stochastic, or when adding additional individuals is not feasible (15, 23).

In a standard case/control study of gene expression with only one sample per individual, typical analysis tests the expression differences between case and control individuals. However, in repeated measures designs with two or more samples per individual, analysis can perform three types of statistical tests: 1) within-individual, 2) a combination of within- and cross-individual, and 3) cross-individual. Within-individual analysis uses a individual-specific baseline to, for example, examine time-course data to identify a differential expression signature of stimulus response, or identify individual-specific expression differences between two cell types each measured in the same set of individuals. In this case, the analysis focuses on the *differences* between the samples from the same individual, and the repeated measures from the same individual capture distinct biology. The simplest form of within-individual analysis is a paired t-test. The combined within- and cross-individual analysis, evaluates within-individual differences between, for example, two time points and then compares the results between cases and controls. Standard RNA-seq software (1, 2, 3, 4, 5, 6) that model these data with fixed effects terms perform well. Alternatively, software using linear mixed models specifically designed for longitudinal analysis of transcriptomic data can be applied (24).

We focus here on cross-individual analysis that considers the *shared* biology of the multiple samples from the same individual. For example, multiple iPSC lines can be generated from the same individual as biological replicates, or multiple brain regions can be assayed from each individual in a case/control study. The repeated measures design can be leveraged to account for biological and technical variability, and model the shared biology of these replicates to increase the statistical power to identify differentially expressed genes. In this case, the multiple replicates from the same individual are considered to be statistically exchangeable after accounting for variation due to covariates such as tissue type, region or technical factors. Since all replicates from the same individual have the same phenotype of interest (i.e. disease status), modeling individual as a fixed effect is not possible because the individual effect is perfectly confounded with the phenotype of interest. Statistically, a fixed effect regression model cannot be fit when the design matrix is not invertible and so parameter estimation and hypothesis testing cannot be performed (25). Therefore, individual must either be modeled as a random effect (22), or the multiple samples from each individual must be collapsed into an individual-level summary.

While summing reads from the multiple samples from the same individual is a simple way to analyze repeated measures data, it has a number of issues. At a basic level it applies equal weight to each individual even though some may have 5 samples while other have only 2. More problematic is the fact that summing reads ignores any biological or technical differences between the multiple samples from the same individual. Some samples may have been processed in a different technical batch, or come from a different tissue or region. Many methods can account for these differences in the full dataset (26, 27, 28), but summing the reads loses this information about within-individual variation.

Moreover, analysts are often interested in tests that allow for heterogeneity of effect sizes within subsets of samples and then perform a joint test of multiple coefficients. For example, in a case/control study of 3 brain regions it may be of interest to allow the case/control effect to vary between brain regions and then perform a joint test of case/control effects with 3 degrees of freedom with an F-test. Obviously, collapsing read counts at the individual level is not compatible with this type of analysis.

Despite the potential of leveraging cross-individual analysis in repeated measures designs, standard differential expression methods do not adequately model the complexity of these repeated measures study designs (1, 2, 3, 4, 5, 6). Recent work has emphasized that applying current methods to cross-individual analysis of repeated measures datasets can result in loss of power or, more problematically, a large number of false positive findings (29, 30). The advantages of repeated measures designs cannot be realized without the proper statistical methods and software.

Statistically, repeated measures data are problematic for existing software because samples from the same individual are correlated, while existing methods assume statistical independence between samples after correcting for covariates. Following our previous work on repeated measures data in genomics (15, 16, 31, 32), this correlation between samples from the same individual can be quantified in terms of the fraction of expression variance explained by variance across individuals. Genes with high variance *across* individuals are expressed at similar levels *within* replicates from the same individual. This departure from statistical independence is substantial and must be considered in any statistical test.

In order to address this issue, Smyth et al. (33) developed the duplicateCorrelation method in the limma workflow (1, 3) to perform differential expression testing while accounting for this correlation structure. Let the correlation between replicates from the same individual for gene *g* be denoted by 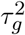. The duplicateCorrelation method uses a single value genome-wide, *τ*^2^, and assumes that the correlation structure for every gene is the same. While this assumption is necessary for dealing with small datasets, current transcriptomic datasets are sufficiently large that this modeling approach is problematic.

Using a single value, *τ*^2^, genome-wide for the correlation within individuals can reduce power and increase the false positive rate in a particular, reproducible way. Consider the correlation value for gene *g*, 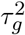, compared to the single genome-wide value, *τ*^2^. When testing a variable that is constant for all replicates of an individual, for genes where 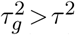, using *τ*^2^ under-corrects for the correlation within individuals so that it increases the false positive rate of gene *g* compared to using 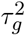. Conversely, for genes where 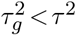, using *τ*^2^ over-corrects for the correlation within individuals so that it decreases power for gene *g*. Increasing sample size does not overcome this issue.

While the use of gene-level random effects has been proposed previously in methodological work, a number of significant hurdles have prevented wider adoption by analysts. Existing methods are either very computational demanding, do not model error structure of RNA-seq data, do not fit easily into existing workflows, or require extensive knowledge of the theory of linear mixed models and details of implementing these models in R. MACAU2 (34) fits a Poisson mixed model for count data for RNA-seq and uses a single random effect to account for multiple samples from the same individual using a pairwise similarity matrix. However, this method does not allow multiple random effects, is not able to fit over-dispersed count models widely used for RNA-seq data (1, 2, 6), and is not scalable to large datasets. Trabzuni, et al. (35) propose a method that fits all of the genes jointly in a linear mixed model, estimates a random effect term modeling the gene by disease interaction, and then considers a genome-wide mixture model of the variance estimates to identify differentially expressed genes. However, this approach does not model count data, is very computationally demanding, and is fit with the commercial package ASReml-R (36). Bryois, et al. (37) apply a linear mixed model to differential expression of RNA-seq data, but do not consider the count-nature of the data and do not provide software.

While a number of generic statistical methods for estimation and hypothesis for testing linear mixed models are available (22, 38, 39), practical application of these methods to RNA-seq data has been limited due to the challenges of 1) implementing these methods for each dataset, 2) high computational cost to fit linear mixed models, 3) directly modeling count data, and 4) uncertainty about the conditions where linear mixed models with gene-level variance terms will outperform existing methods.

Here we present a statistical software package, dream (<u>d</u>ifferential expression for <u>re</u>pe<u>a</u>ted <u>m</u>easures), that addresses each of these hurdles, and outperforms existing methods for cross-individual tests in repeated measures datasets.

## MATERIALS AND METHODS

Linear mixed models are commonly applied in biostatistics in order to account for the correlation between observations from the same individual in repeated measures studies (22, 40). We start with a description of a simple linear model for differential expression analysis and build towards the dream model.

### Linear models for differential expression

Consider a linear model for a single gene

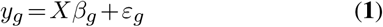

where *y*_*g*_ is a vector of log_2_ counts per million for gene *g*, the matrix *X* stores covariates as columns, *β*_*g*_ is the vector of regression coefficients, and *ε*_*g*_ is normally distributed error. In order to account for heteroskedastic error from RNA-seq counts, the error takes the form

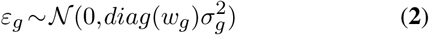

where 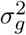 is the residual variance, and *w*_*g*_ is a vector of precision weights (1). Precision weights can be learned from the data in order to account for counting error in RNA-seq or variation in sample quality (1, 41). In this case, the estimates 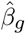 can be obtained by a closed form least squares model fit. Hypothesis testing is performed by specifying a contrast matrix *L* that is a linear combination of the estimated coefficients, and evaluating the null model

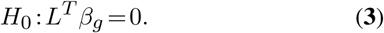

Alternatively, an F-test jointly testing multiple coefficients can be applied.

### Accounting for repeated measures with a two step model: duplicateCorrelation

The most widely used approach for handing repeated measures in differential expression analysis is the `duplicateCorrelation()` function available in limma (3). This approach involves two steps. In the first step, a linear mixed model is fit for each gene separately, and only allows a single random effect. The model is

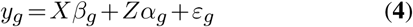

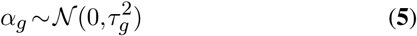

where *Z* is the design matrix for the random effect, with coefficients *α*_*g*_ drawn from a normal distribution with variance 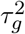. After fitting this model for each gene, a single genome-wide variance term is computed according to

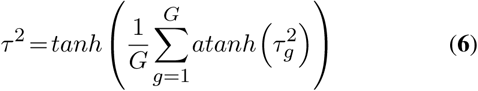

where *G* is the number of genes, *tanh* is the hyperbolic tangent and *atanh* is its inverse.

In the next step, this single variance term, *τ*^2^, is then used in a generalized least squares model fit for each gene, blocking by individual:

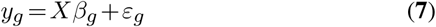

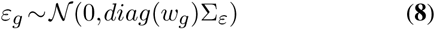

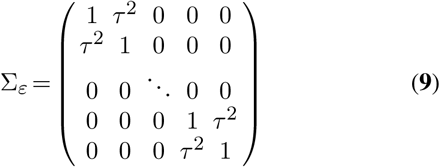

where Σ_*ε*_ is the covariance between samples and considers the correlation between samples from the same individual. Note that the same *τ*^2^ value is used for all genes.

The duplicateCorrelation method allows the user to specify a single random effect usually corresponding to donor. So it can’t model multi-level design. Moreover, duplicateCorrelation estimates a single variance term genome-wide even though the donor contribution of a particular gene can vary substantially from the genome-wide trend (31). Using a single value genome-wide for the within-donor variance can reduce power and increase the false positive rate in a particular, reproducible way as described in the introduction.

Using the single variance term genome-wide and using the *tanh* and *atanh* are designed to address the high estimation uncertainly for small gene expression experiments. However, using this single variance term has distinct limitations. First, it ignores the fact that the contribution of the random effect often varies widely from gene to gene (31). Using a single variance term to account for the correlation between samples from the same individuals over-corrects for this correlation for some genes and under-corrects for others. In addition, it is a two step approach that first estimates the variance term and then estimates the regression coefficients. Thus, it does not account for the statistical uncertainty in the estimate of *τ*^2^. Finally, it does not account for the fact that estimating the variance component changes the null distribution of 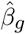. Specifically, estimating variance components in a linear mixed model can substantially change the degrees of freedom of the distribution used to approximate the null distribution for fixed effect coefficients (39, 42, 43, 44, 45). Ignoring this issue can lead to false positive differentially expressed genes.

### Dream model

The dream model extends the previous model in order to

- enable multiple random effects
- enable the variance terms to vary across genes
- approximate degrees of freedom of hypothesis test for each gene and contrast from the data in order to reduce false positives

The definition of the dream model follows directly from the definition of the previous models. First, consider a linear mixed model for gene *g* with an arbitrary number of random effects:

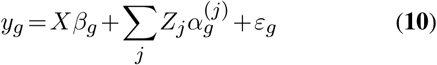

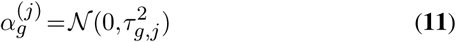

where *Z*_*j*_ is the design matrix for the *j*^*th*^ random effect, with coefficients 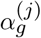 drawn from a normal distribution with variance 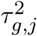. As before, heteroskedastic errors are modeled with precision weights with

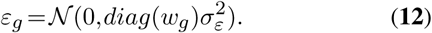

In this case, estimates of coefficients 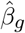 and variance Components 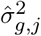 must be obtained via an iterative optimization algorithm (38).

For the linear model and generalized least squares model described above, the degrees of freedom of the hypothesis test is fixed based on the number of covariates and the sample size:

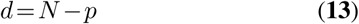

where *N* is the number of samples, and *p* is the number of covariates. For the linear mixed model, we can explicitly account for the fact that estimating the random effect changes the degrees of freedom of the distribution used to approximate the null distribution (39, 42, 43, 44, 45). Thus let *d*_*g,C,L*_ be the degrees of freedom of the hypothesis test for gene *g* for the specified set of covariates, *C*, and contrast matrix, *L*. We omit the statistical details here, but *d*_*g,C,L*_ can be estimated from the model fit using the very fast Satterthwaite approximation (39, 44) (the default in dream) or the more accurate but computationally demanding Kenward-Roger approximation (43, 45) used by dream-KR.

### Modeling measurement error in RNA-seq counts

The limma package models measurement error in the RNA-seq counts by estimating precision weights for each observation (1). The `voom()` function does this by fitting a smooth function to the square root residual standard deviation regressed on the log_2_ counts. However, `voom()` can only model variables as fixed effects, and so cannot consider within-individual variation in estimating the precision weights. In `voomWithDreamWeights()`, we apply an identical procedure except that the residuals are computed by fitting a linear mixed model specified by the user. A dream analysis can use precision weights computed by either `voom()` or `voomWithDreamWeights()`.

### Features of dream method

Dream enables powerful analysis of repeated measures data while properly controlling the false positive rate. Dream combines:

- random effects estimated separately for each gene (31)
- ability to model multiple random effects (38)
- fast hypothesis testing for fixed effects in linear mixed models (39), including:
  **–** tests of single coefficients
  **–** linear contrasts specifying a linear combination of coefficients
  **–** joint hypothesis testing of multiple coefficients using an F-test
- small sample size hypothesis test to increase power (45)
- precision weights to model measurement error in RNA-seq counts (1, 41)
- seamless integration with the widely used workflow of limma (3)
- parallel processing on multi-core machines (46)

### Software

The dream method is available in the `dream()` function in the variancePartition (31) package (http://bioconductor.org/packages/variancePartition) from Bioconductor version ≥ 3.7.

### Implementation

Linear mixed models are estimated using the lme4 package (38). Estimating the residual degrees of freedom is performed with either Satterthwaite approximation (44) in the lmerTest package (39) or the Kenward-Roger approximation (43) in the pbkrtest package (45). Parallel processing of thousands of genes on a multi-core computer is performed with BiocParallel (46). Visualization is performed with ggplot2 (47).

### Simulations

The true expression values were simulated from a linear mixed model with 4 components: 1) variance across individuals; 2) variance across two disease classes (i.e. cases versus controls); 3) variance across two batches; and 4) residual variance. All samples from a given individual have the same disease status, and analysis considers the cross-individual test of differential expression between cases and controls. Samples are randomly assigned to one of two batches. Including a batch component here models the expression heterogeneity across technical batches, but also across two brain regions, or tissues types as is common in real data. For each gene, the variance fractions for these 4 components were sampled from a beta distribution with parameters set to give a specified mean and variance described below. The residual variance was set so that the variance fractions summed to 1. If the randomly draw values sum to more than one, the residual variance is set to 0.05 and the other components are scaled accordingly. The variance component values were based on examining the variance fractions estimated with variancePartition across many datasets (15, 16, 31, 32). In each simulation, 500 genes were randomly selected to be differentially expressed between cases and controls, and for all other genes the disease component was set to zero.

At the implementation level, simulating expression data is divided into three steps: 1) Simulate relative expression values, 2) convert these into multiplicative fold changes compared to a baseline, 3) simulate RNA-seq counts from negative binomial model given the expected multiplicative fold change values.

Let *y*_*j*_ be the vector of *relative* expression values for gene *j* across all individuals that varies from −∞ to ∞. Let *η*_*k*_ correspond to the *k*^*th*^ component where *k* ∈ (Individual, Disease, Batch, Noise) so that, for example, *η*_Individual_ represents the vector of expected expression attributable to variance across individuals. Since each variable represented by *k* has a different number of levels, let 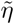 be the standardized version of *η* with a mean of 0 and variance 1. The expression value of gene *j* is simulated according to

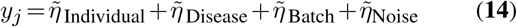

where each variance component *η*_*k*_ is drawn according to

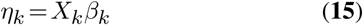

where *X*_*k*_ is the matrix of ANOVA coded indicator values for variable *k*. The number of columns in *X*_*k*_ equals the number of categories in variable *k* so that Disease has two categories, Batch has 2 categories and Individual has *N* categories. The vector *β*_*k*_ gives the expected value for each of the categories in variable *k*. These expected values are drawn from a normal distribution according to

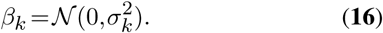

Since the expression values are simulated based on standardized variance components, 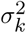 corresponds to the fraction of variance attributable to component *k*. Since fractions naturally fall between 0 and 1, they can be drawn from a beta distribution. The beta distribution is usually parameterized in terms of two shape parameters, *α* and *β*, but here we parameterize it with a mean and variance by matching the moments of the distribution. The simulated mean for each component corresponds to the expected variance fractions and were motivated based on variancePartition across many datasets (15, 16, 31, 32). The simulated variance fractions are draw according to

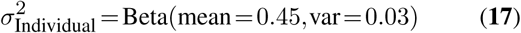

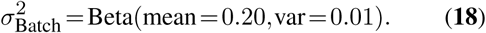

For the 500 genes where disease has an effect on expression,

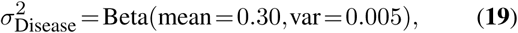

and the value of 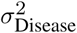 is subtracted from 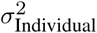, since all samples from the same individual have the same disease status. For all other genes, 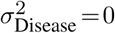.

The random noise component is drawn so that the variance fractions sum to 1, but is set to a minimum of 0.05:

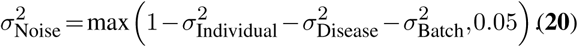

Given, *y*_*j*_, the vector of relative expression values for gene *j*, convert them into multiplicative fold change values with a minimum of 1 according to

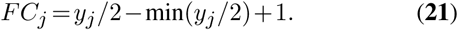

These simulated multiplicative fold change values are passed to polyester v1.14.0 (48) to produce counts with biologically realistic negative binomial error. The number of reads per gene is selected based on gene length using the ‘mean model’ from polyester. Otherwise, polyester defaults were used.

Expression values were simulated following above procedure for 20,738 protein coding genes from GENCODE v19. Approximately 57 million total reads counts were generated for each sample (mean: 57.3M, median: 55.0M, sd: 24.4M). Simulations using a range of values for each of these parameters did not change the conclusions. Simulation and data analysis code is available at https://github.com/GabrielHoffman/dream_analysis.

### Data analysis

Data for Timothy syndrome (20) was downloaded from GEO at GSE25542. Data for childhood onset schizophrenia (15) was downloaded from https://www.synapse.org/#!Synapse:syn9907463. Post mortem brain RNA-seq data from Alzheimer’s and controls (10) was downloaded from https://www.synapse.org/#!Synapse:syn3159438. Analysis was performed on individuals from European ancestry that were assayed in each of 4 brain regions (Brodmann areas 10, 22, 36 and 4), had ApoE genotype data, had Braak stage information, and were either controls or definite AD patients (i.e. possible and probable cases were excluded). Differential expression analysis corrected for batch, sex, RIN, rRNA rate, post mortem interval, mapping rate and ApoE genotype.

Data from GENESIPS (16) was obtained from GEO at GSE79636. Data from Warren, et al. (17) was obtained from GEO at GSE90749. Data from Mariani, et al. (19) was obtained from recount2 (49) at SRP047194. Enrichment analysis was performed with cameraPR (50) in the limma package (3). In order to avoid using arbitrary cutoffs to identify differentially expressed genes, gene set enrichments were evaluated by applying cameraPR to the differential expression test statistics from each analysis. The fraction of expression variation explainable by cis regulatory variants was obtained from Gamazon et al. (51) and Huckins, et al. (52). Reproducible analysis code, figures, and statistics from differential expression and enrichment analyses are available at https://github.com/GabrielHoffman/dream_analysis.

## RESULTS

### Biologically realistic simulations demonstrate performance of dream

The performance of dream was compared to current methods on biologically realistic simulations. The methods can be divided into six categories: 1) dream using default settings or a Kenward-Roger approximation (termed dream-KR, see Methods), which is more powerful but much more computationally demanding; 2) duplicateCorrelation from the limma/voom workflow (3); 3) MACAU2 (34); in addition to DESeq2 (2) and limma/voom(1) applied to: 4) including all samples but ignoring the repeated measures design; 5) with only a single replicate per individual; 6) summing the reads across biological replicates from the same individual.

The two dream methods are more powerful than the other methods (**Figure 1**). Across a range of simulations of 4 to 50 individuals each with 2 to 4 biological replicates, the two dream methods have a lower false discovery rate (**Figure 1A, S1**), better precision-recall curves (**Figure 1B, S2**), and larger area under the precision-recall (AUPR) curve (**Figure 1C, S3**). A test of differential expression must control the false positive rate accurately in order to be useful in practice. As expected (29, 30), the methods that include all samples but ignore the correlation structure do not control the false positive rate (**Figure 1D**). Importantly, analysis that sums reads from multiple replicates controls the false positive rate as expected, and has better power than using only a single replicate. Yet summing has lower power than methods that model the correlation structure of the full dataset.

**Figure 1.**
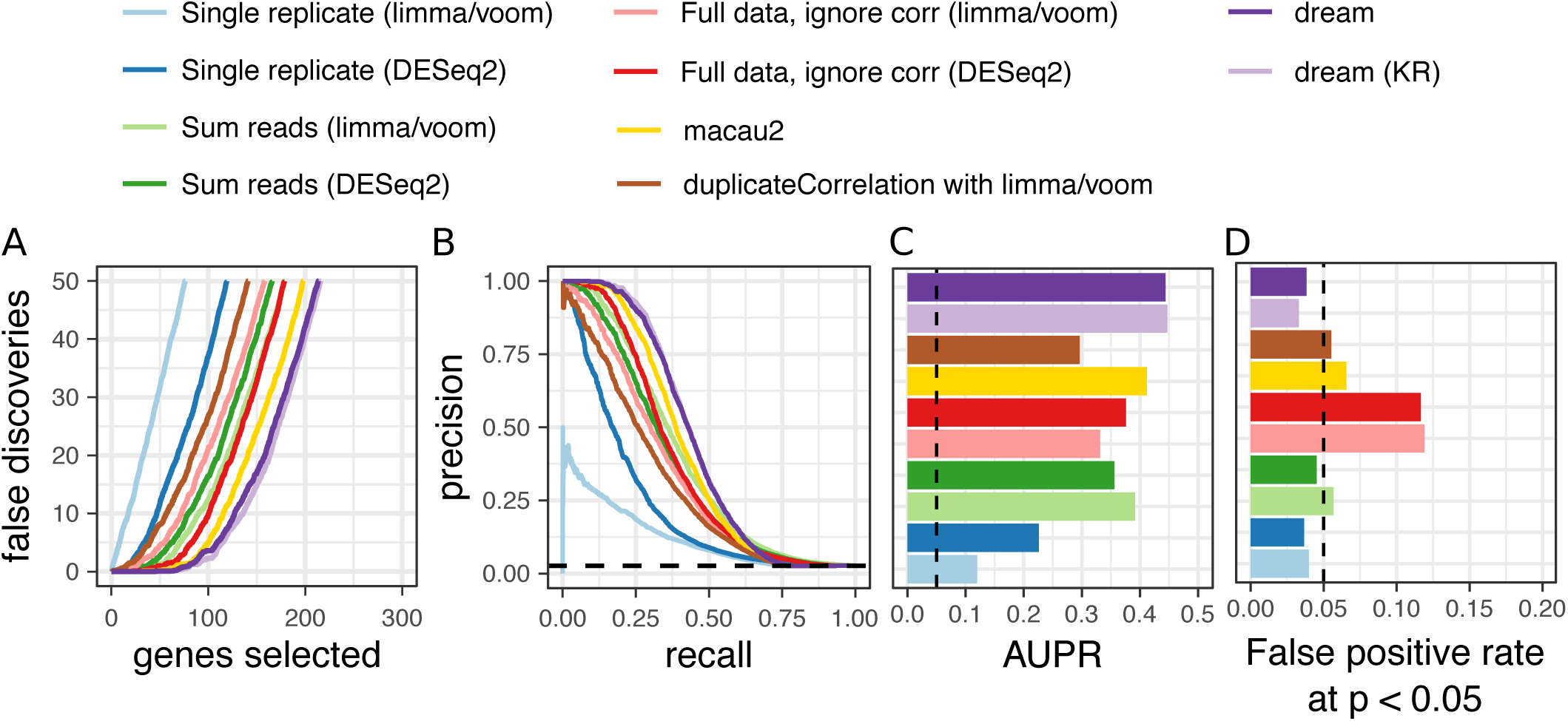
Performance on biologically realistic simulated data. **A,B,C,D)** Performance from 50 simulations of RNA-seq datasets of 8 individuals each with 3 replicates. **A)** False discoveries plotted against the number of genes called differentially expressed by each method. **B)** Precision-recall curve showing performance in identifying true differentially expressed genes. Dashed line indicates performance of a random classifier. **C)** Area under the precision-recall (AUPR) curves from **(B)**. Dashed line indicates AUPR of a random classifier. **D)** False positive rate at p *<* 0.05 evaluated under a null model were no genes are differentially expressed illustrates calibration of type I error from each method. As indicated by the dashed line, a well calibrated method should give p-values *<* 0.05 for 5% of tests under a null model.

Aggregating results across many simulation conditions reveals trends as the number of individuals and replicates increases (**Figure 2**). The lack of type I error control for methods that ignore the correlation structure, as well as MACAU2, is present in all simulation conditions (**Figure 2A, S4**). Even more concerning, increasing the number of repeated measures can dramatically increase the false positive rate. Notably, duplicateCorrelation shows a slight increase in type I error at larger sample sizes. For MACAU2, the type I error is very inflated for small samples sizes but decreases for larger datasets. Higher type I error can translate into hundreds of false positive differentially expressed genes even when no genes are truly differently expressed (**Figure 2B**). Importantly, both versions of dream accurately control the type I error with sufficient sample size.

**Figure 2.**
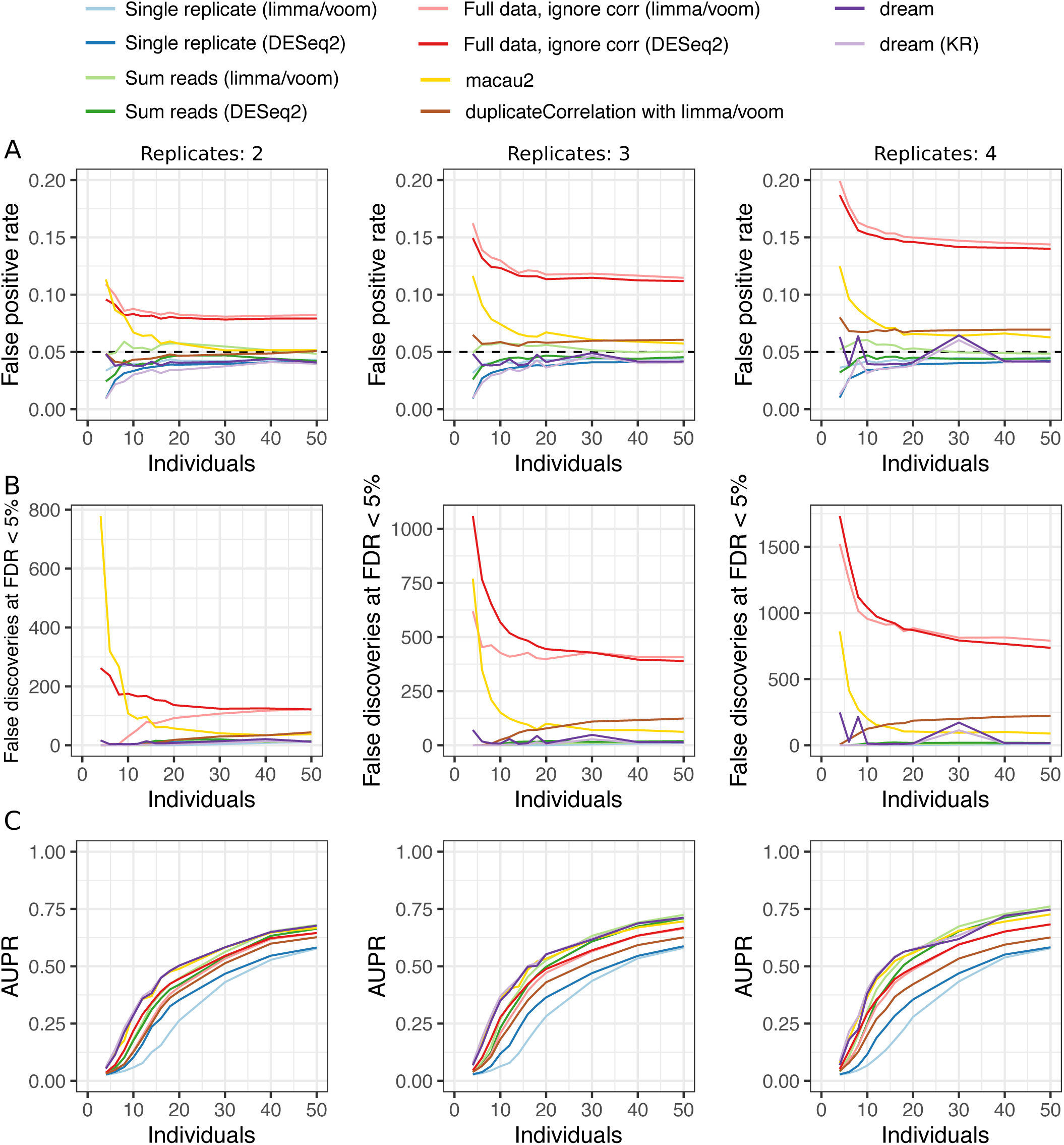
Performance summary for simulations with a range of individuals and replicates. Simulations were performed on 4 to 50 individuals with between 2 to 4 replicates. For each condition, 50 simulations were performed for a total of 1800. **A)** False positive rate at p *<* 0.05 for simulations versus the number of individuals and replicates. Black dashed line indicates target type I error rate of 0.05. **B)** Number of genes passing FDR cutoff of 5% under the null simulations. Values shown are averaged across 50 simulations. **C)** AUPR for simulations versus the number of individuals and replicates.

The two versions of dream give the highest AUPR across all simulation conditions (**Figure 2C**) while properly controlling the false positive rate. In addition, MACAU2 also produces a competitive AUPR, but lack of type I error control and high computational cost is problematic (**Figure S5A**). While dream-KR gives the best performance, especially at small sample sizes, the computational time required can be prohibitive. Using dream with the default settings performs nearly as well in simulations, but can be 2-20x faster (**Figure S5B**). For datasets with greater than 500 individuals, dream is also 5-10x faster than duplicateCorrelation.

### Analysis of expression profiling datasets with dream gives biological insight

Applying dream to empirical data gives biological insight for 3 neuropsychiatric diseases with different genetic architectures. In order to avoid using arbitrary p-value or FDR cutoffs to identify differentially expressed genes, gene set enrichments were evaluated using cameraPR (3, 50) to compare the differential expression test statistics from genes in a given gene set to the genome-wide test statistics.

Alzheimer’s disease is a common neurodegenerative disorder with a complex genetic architecture (53)(**Figure 3**). In analysis of RNA-seq data from 4 regions of post mortem brains from 26 individuals (10), dream identified known patterns of dysregulation in genes involved in adipogenesis, inflammation and monocyte response associated with Braak stage, a neuropathological metric of disease progression (**Figure 3A, S6**). Here we allow the disease effect to be different in each brain region and then test if the sum of the 4 coefficients is significantly different from zero using a linear contrast. Applying duplicateCorrelation only recovered a subset of these findings and produced larger false discovery rates across many biologically relevant gene sets. Notably, the difference between dream and duplicateCorrelation is due to the way that these methods account for expression variation explained by variance across individuals (**Figure 3B**). Genes with correlation within individuals, 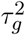, that is larger than the genome-wide average, *τ*^2^, are susceptible to being called as false positive differentially expressed genes by duplicateCorrelation. Conversely if 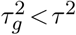, then dream will tend to give a more significant p-value than duplicateCorrelation. For example, consider TUBB2B where 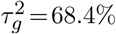 compared to the genome-wide *τ*^2^ = 38.8%. Here, duplicateCorrelation under-corrects for the correlation structure and gives a p-value of 1.3e-10 while dream uses a gene-specific correlation to give p-value of only 7.7e-4 (**Figure 3C**). Finally, we note that performing a joint F-test of these coefficients with with 4 degrees of freedom gives similar results (**Figure S7**).

**Figure 3.**
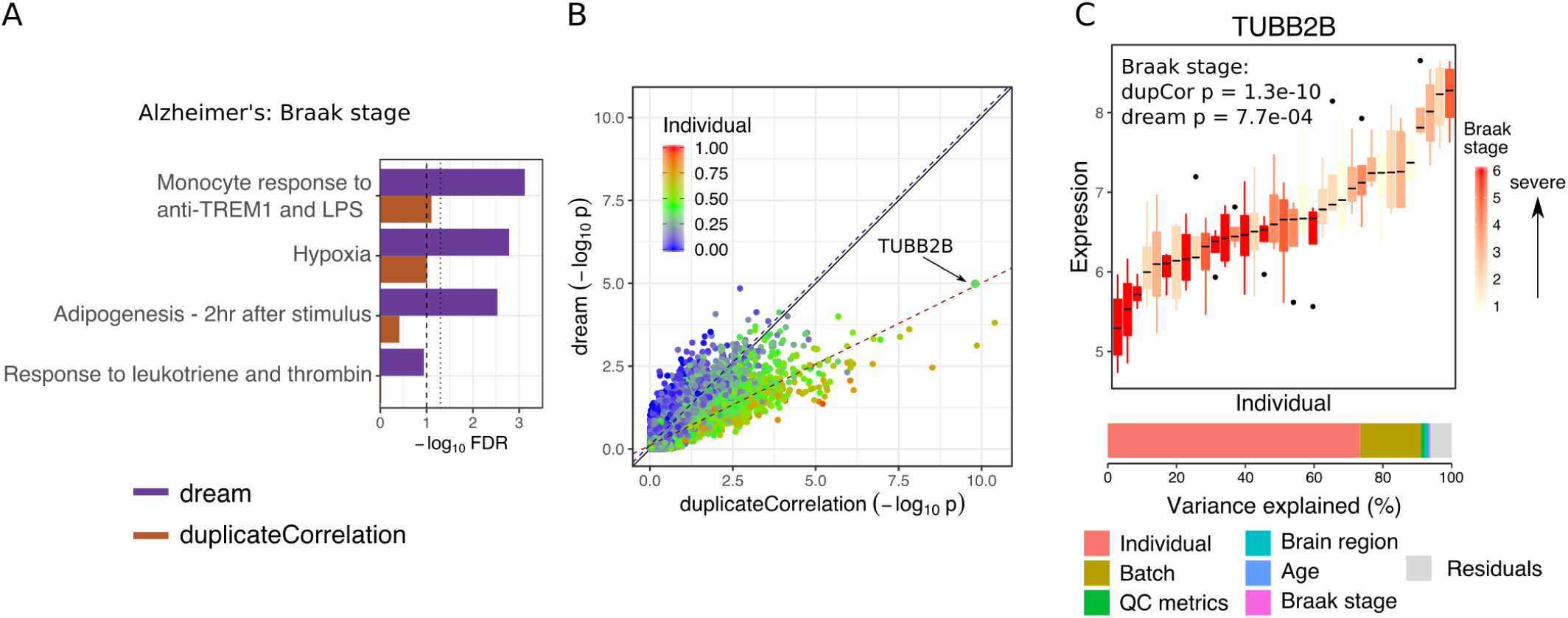
Application to transcriptome data from Alzheimer’s disease. **A**) Gene set enrichment FDR for genes associated with Braak stage. Results are shown for dream and duplicateCorrelation. Lines with broad and narrow dashes indicate 10% and 5% FDR cutoff, respectively. **B**) Comparison of −log_10_ p-values from applying dream and duplicateCorrelation to Braak stage. Each point is a gene, and is colored by the fraction of expression variation explained by variance across individuals. Black solid line indicates a slope of 1. Dashed line indicates the best fit line for the 20% of genes with the highest (red) and lowest (blue) expression variation explained by variance across individuals. **C**) Expression of TUBB2B stratified by individual and colored by Braak stage so that each box represents the expression in the multiple samples from a given individual. Bar plot of variance decomposition shows that 68.4% of variance is explained by expression variance across individuals. Since this value is much larger than the genome-wide mean, duplicateCorrelation under-corrects for the repeated measures.

Childhood onset schizophrenia is a severe neurodevelopment disorder, but the genetic cause is complex with patients having a higher rate of schizophrenia-associated copy number variants, as well as higher schizophrenia polygenic risk scores (54, 55). RNA-seq data was generated from iPSC-derived neurons and neural progenitor cells from 11 patients with childhood onset schizophrenia (15) and 11 controls with up to 3 lines per donor and cell type. Differential expression analysis was performed in each cell type (**Figure 4**). The relationship between results from dream and duplicateCorrelation is again well captured by the correlation across individuals at the gene level (**Figure 4A**). Analysis with dream identified gene sets involved in neuronal function at the 5% and 10% FDR levels that were not identified by duplicateCorrelation (**Figure 4B, S8**).

**Figure 4.**
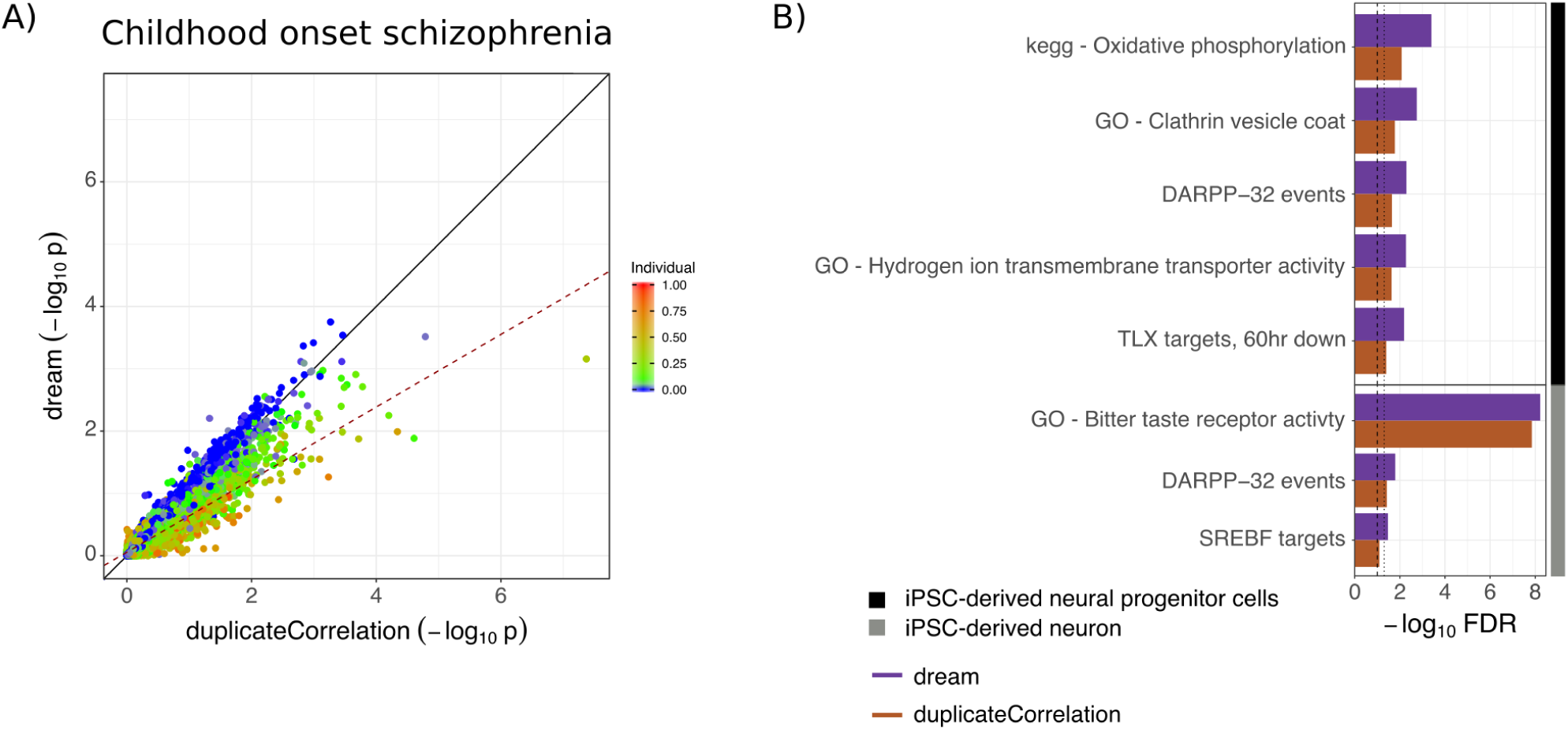
Application to transcriptome data from childhood onset schizophrenia. **A**) Comparison of −log_10_ p-values from applying dream and duplicateCorrelation to disease status in neurons. Each point is a gene, and is colored by the fraction of expression variation explained by variance across individuals. Black solid line indicates a slope of 1. Dashed line indicates the best fit line for the 20% of genes with the highest (red) and lowest (blue) expression variation explained by variance across individuals. **B**) Gene set enrichment FDR for genes associated with disease status in iPSC-derived neurons and neural progenitor cells. Results are shown for dream and duplicateCorrelation. Lines with broad and narrow dashes indicate 5% and 10% FDR cutoff, respectively.

Timothy syndrome is a monogenic neurodevelopmental disorder caused by variants in the calcium channel CACNA1C. Induced pluripotent and derived cell types were generated from 2 affected and 4 unaffected individuals an expression was assayed by microarray (20, 56). Since up to 6 lines were generated per donor for each cell type, it is necessary to account for the repeated measures design. Analysis with dream removed many differentially expressed genes identified by duplicateCorrelation where the signal was driven by variation across individual rather than variance across disease status (**Figure S9**).

### False positives driven by genetic regulation

Since the relationship between results from dream and duplicateCorrelation depend on 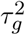 and *τ*^2^, we examined which genes had large 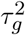 values and were thus highly susceptible to being called as a false positive differentially expressed gene by duplicateCorrelation. Since 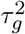 is a metric of the expression variation across individuals, we hypothesized that this variation was driven by genetic regulation of gene expression. In order to test this, we use large-scale RNA-seq datasets from the post mortem human brains from the CommonMind Consortium (57) and whole blood from Depression Genes and Networks (58) where eQTL analysis had already been performed. Transcriptome imputation was performed by training an elastic net predictor for each gene expression trait using only cis variants (51, 52). For each gene, the fraction of expression variation explainable by cis regulatory variants was termed ‘eQTL R^2^’.

Comparing the eQTL R^2^ of each gene to the −log_10_ p-values from differential expression using dream and duplicateCorrelation revealed a striking trend (**Figure 5, S10**).

**Figure 5.**
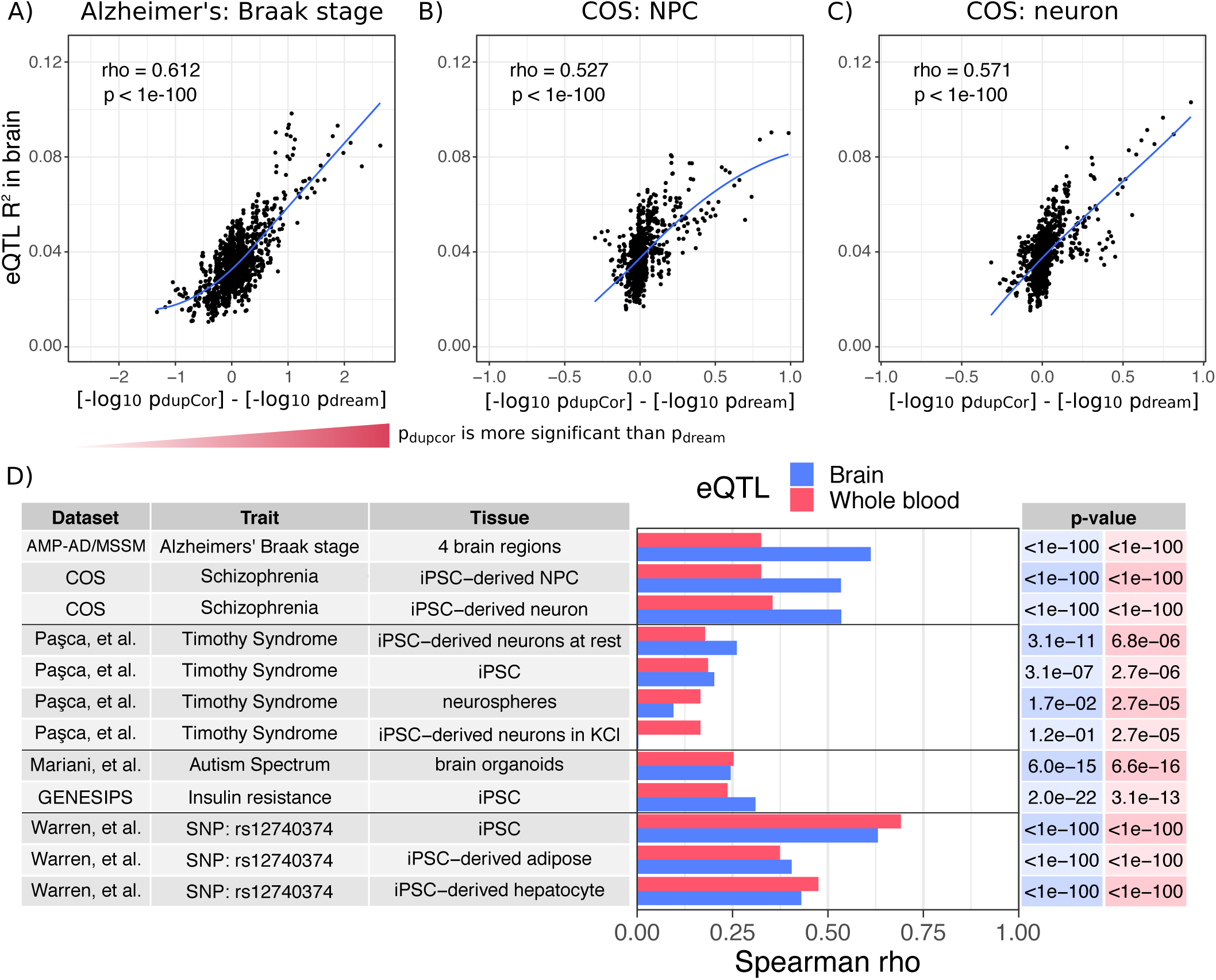
Genes falsely called differentially expressed tend to be under strong genetic regulation. For each gene the fraction of expression variation explainable by cis-eQTLs is compared to the difference in −log_10_ p-value from duplicateCorrelation and dream differential expression analysis. Due to the large number of genes, a sliding window analysis of 100 genes with an overlap of 20 was used to summarize the results. **A,B,C**) For each window, the average fraction of expression variation explainable by cis-eQTLs (i.e. eQTL R^2^) in the CommonMind Consortium (57) and average difference in −log_10_ p-values from the two methods are reported when differential expression analysis is performed on **A**) Alzheimer’s Braak stage from post mortem brains, and schizophrenia status from iPSC-derived neural progenitor cells and **C**) iPSC-derived forebrain neurons. Spearman rho correlations and p-values are shown along with loess curve. **D**) Summary of Spearman rho correlations between eQTL R^2^ and the difference between −log_10_ p-value from duplicateCorrelation and dream for 12 analyses in 6 datasets. P-values for each correlation are shown on the right in the corresponding color. Results are shown for eQTL R^2^ from brains from the CommonMind Consortium (57) and whole blood from Depression Genes and Networks (DGN) dataset (58). Note that differential expression analysis compared disease to control individuals in each tissue from each dataset, except for AMP-AD/MSSM where Alzheimer’s Braak stage is a quantitative metric, and Warren et al. (17) where the variable of interest was the SNP rs12740374.

For differential expression analysis with Braak stage from the Alzheimer’s dataset, genes that were more significantly differentially expressed by duplicateCorrelation compared to dream had a much higher expression variation explainable by cis-eQTLs in post mortem brain (**Figure 5A**). This strong trend was also seen in differential expression results for childhood onset schizophrenia in neural progenitor cells (**Figure 5B**) and neurons (**Figure 5C**).

To see how widespread this trend was, we performed 12 differential expression analysis from 6 independent expression datasets. Comparing the differential expression results with the eQTL R^2^ from brain and whole blood showed correlations that were highly significant in all datasets, but more importantly, that were surprisingly large. While the correlation with eQTL R^2^ from brain was larger in most cases because we considered many brain and neuronal datasets, the signal from whole blood was still very robust. Moreover, expression datasets from iPSC, iPSC-derived adipocytes and iPSC-derived hepatocytes showed the same trend.

The analysis of these last 3 cell types from Warren et al. (17) is notable because the cohort was designed to have an equal number of individuals who were homozygous reference as homozygous alternate at rs12740374, a SNP associated with cardiometabolic disease. Even though the variable used in the differential expression analysis was itself the allelic state at this SNP, the results from the duplicateCorrelation analysis compared to dream were still strongly correlated with eQTL R^2^. Thus this trend is independent of the biology of cell type and trait of interest, but is instead driven by genetic regulation of gene expression.

## DISCUSSION

As study designs for transcriptome profiling experiments becomes more complex (7, 8, 9, 10, 11, 12, 13, 14, 15, 16, 17, 18, 19, 20), proper statistical methods must be used in order to take full advantage of the power of these new datasets and, more importantly, protect against false positive findings. The results of our biologically realistic simulation study indicate that analyzing the full repeated measures dataset while properly accounting for the correlation structure gives the best performance for identifying differential expression, compared with methods that omit samples, use individual-level summaries or ignore the correlation structure.

We have demonstrated that dream has superior performance in biologically realistic simulations while retaining control of the false positive rate. Furthermore, dream is able to identify biologically meaningful gene set enrichments in two neuropsychiatric disorders with different genetic architectures where the current standard for repeated measures designs in transcriptomics, duplicateCorrelation, cannot.

Relating the performance of dream and duplicateCorrelation to the expression variation across individuals at the gene level gives a first principles framework for understanding the empirical behavior of these methods. Based on this understanding, we observe how genes with expression variation across individuals *below* the genome-wide mean benefit from increased power using dream. Meanwhile, genes with expression variation across individuals *above* genome-wide mean benefit from proper control of the false positive rate compared to duplicateCorrelation.

We further demonstrated how genes under strong genetic regulation are be particularly susceptible to being called as false positives by differential expression analysis with duplicateCorrelation. Since this effect is attributable to strong eQTL’s, these differential expression results can be reproducible across multiple datasets despite being false positive findings unrelated to the biological trait of interest. Notably, dream uses a gene-specific variance term 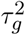 and so it is not susceptible to these artifactual findings.

Here we focus on cross-individual analysis of repeated measures design because of the limitations of existing statistical software for this application, and the concern about false positive findings raised by recent work (29, 30). Analysis of repeated measures data is a broad field (22, 40) that includes with-individual and combined cross- and within-individual tests. Existing statistical methods for RNA-seq data perform well on those applications (1, 2, 3, 4, 5, 6).

## CONCLUSION

Since dream is built on top of the limma (3) and variancePartition (31) workflow, it can easily accommodate expression quantifications from multiple software packages including featureCounts (59), kallisto (60), salmon (61), and RSEM (62), among others. Moreover, dream works seamlessly for differential analysis of ATAC-seq or histone modification ChIP-seq data. Finally, with scaleable single cell RNA-seq on the horizon, future studies will need to perform differential expression analysis with thousands of cells (i.e. repeated measures) from each individual. (11, 63). The power, type I error control, simple R interface, speed and flexibility of dream enables analysis of transcriptome and functional genomics data with repeated measures designs.

## ACKNOWLEDGEMENTS

We thank Laura Huckins for providing the eQTL R^2^ values. This work was supported by NIMH grants U01MH116442, R01MH109677, R01MH109897, R01MH110921, NIA grant R01AG050986, and Veterans Affairs merit grant BX002395 to P.R. G.E.H. is partially supported by a NARSAD Young Investigator Award 26313 from the Brain and Behavior Research Foundation. This work was supported in part through the computational resources and staff expertise provided by Scientific Computing at the Icahn School of Medicine at Mount Sinai.

---

**Figure S 1.**
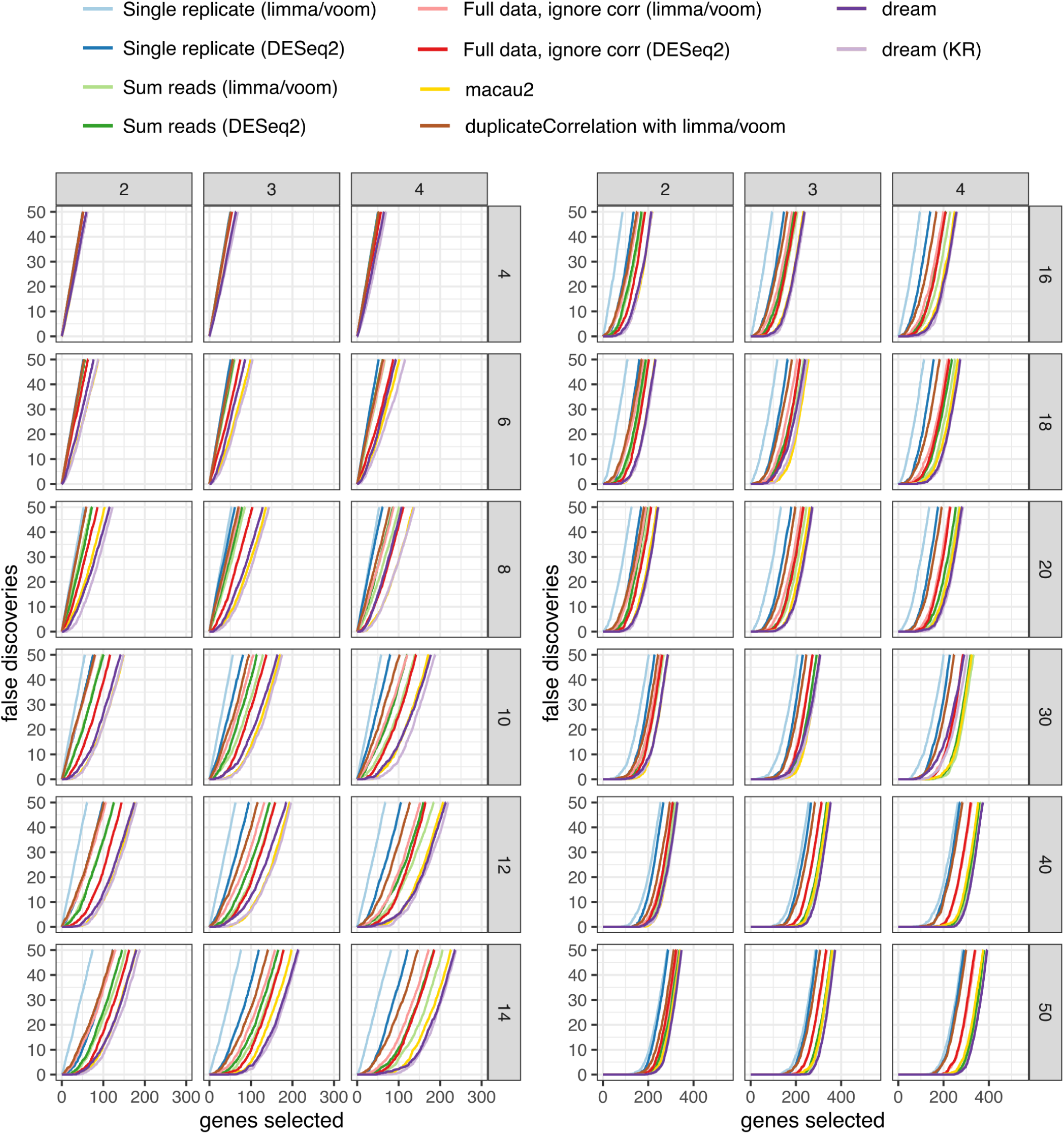
False discovery rates for multiple simulation conditions. False discoveries plotted against the number of genes called differentially expressed by each method. Results are shown for between 4 and 50 individuals (rows) and 2 to 4 replicates (columns). For each combination, 50 simulated datasets were analyzed.

**Figure S2.**
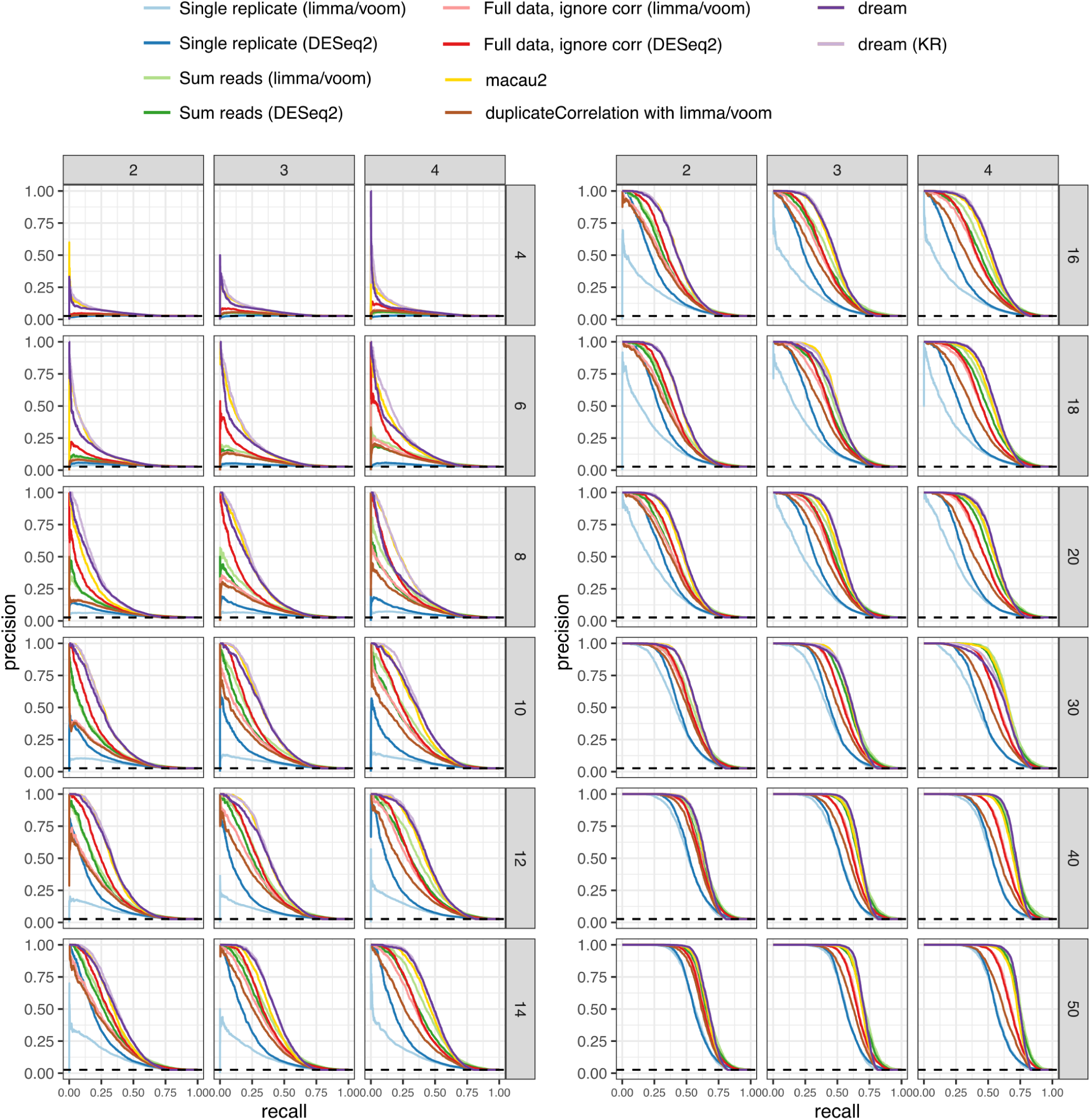
Precision-recall curves for multiple simulation conditions. Plots shows performance in identifying true differentially expressed genes. Dashed lined indicates performance of a random classifier. Results are shown for between 4 and 50 individuals (rows) and 2 to 4 replicates (columns). For each combination, 50 simulated datasets were analyzed.

**Figure S 3.**
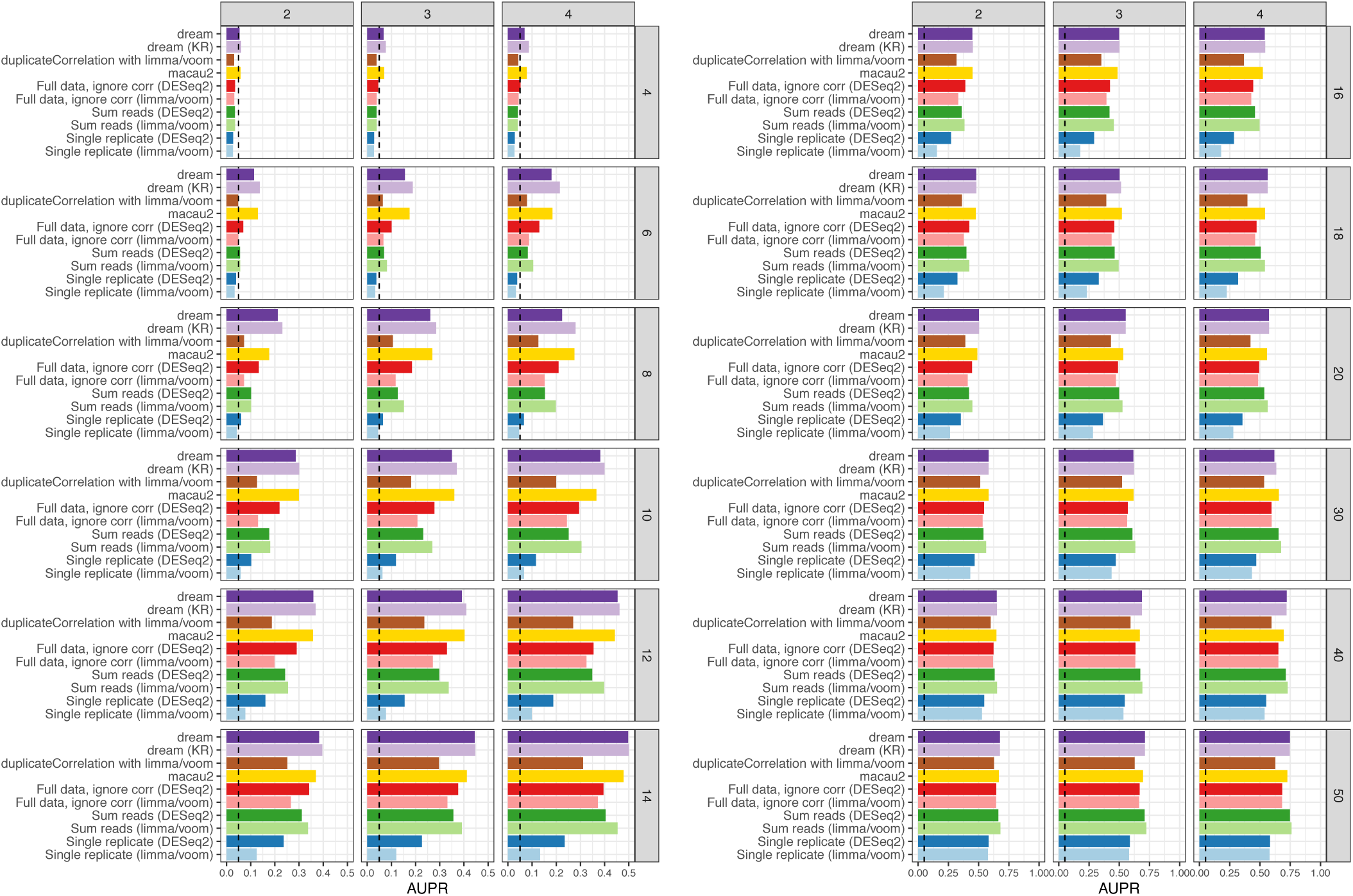
Area under the precision-recall (AUPR) for multiple simulation conditions. Dashed line indicates AUPR of a random classifier. Results are shown for between 4 and 50 individuals (rows) and 2 to 4 replicates (columns). For each combination, 50 simulated datasets were analyzed.

**Figure S4.**
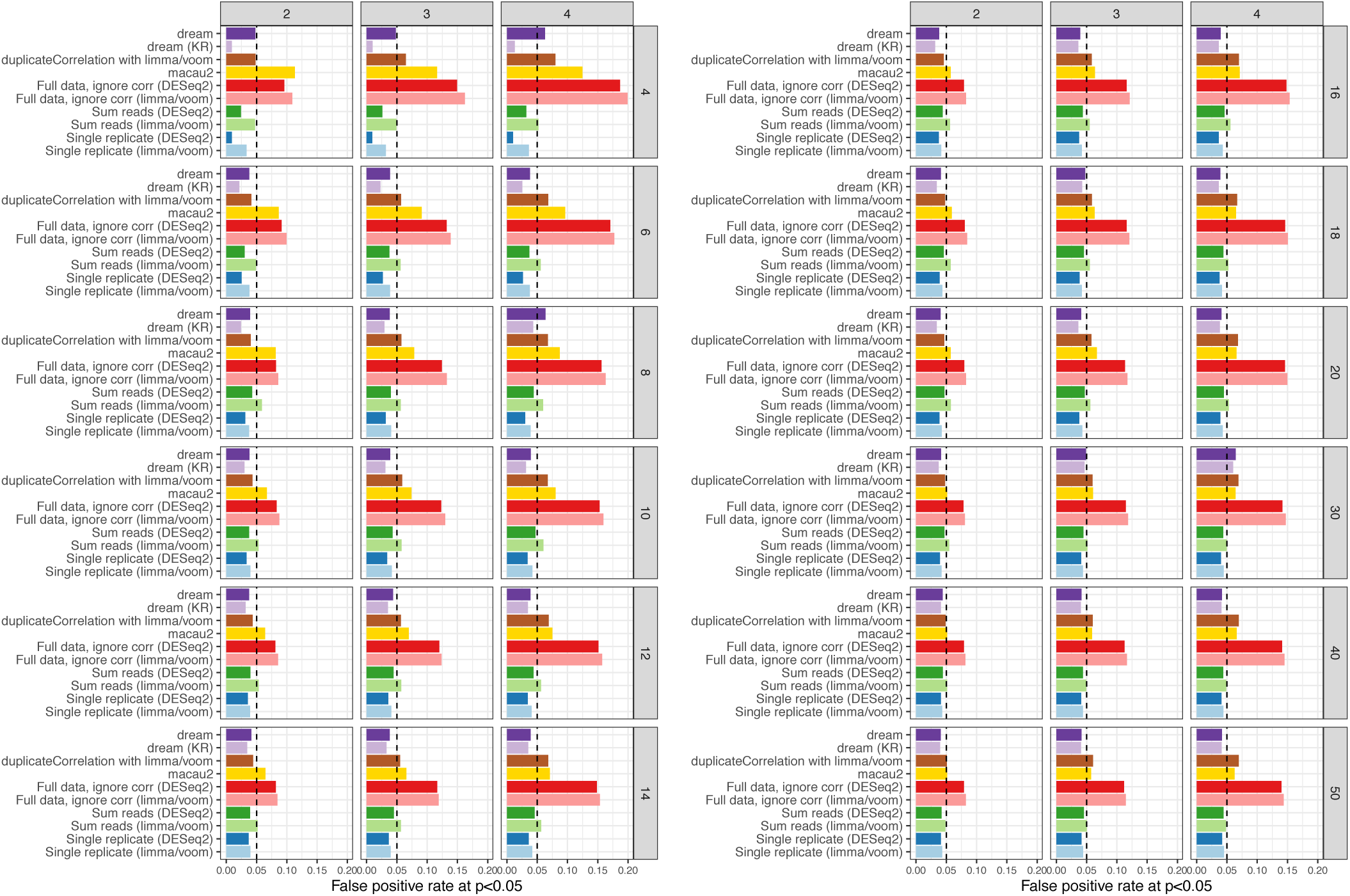
False positive rate for multiple simulation conditions. False positive rate at p *<* 0.05 evaluated under a null model were no genes are differentially expressed illustrates calibration of type I error from each method. As indicated by the dashed line, a well calibrated method should give p-values *<* 0.05 for 5% of tests under a null model. Results are shown for number of individuals between 4 and 50 (rows), and replicates between 2 and 4 (columns).

**Figure S 5.**
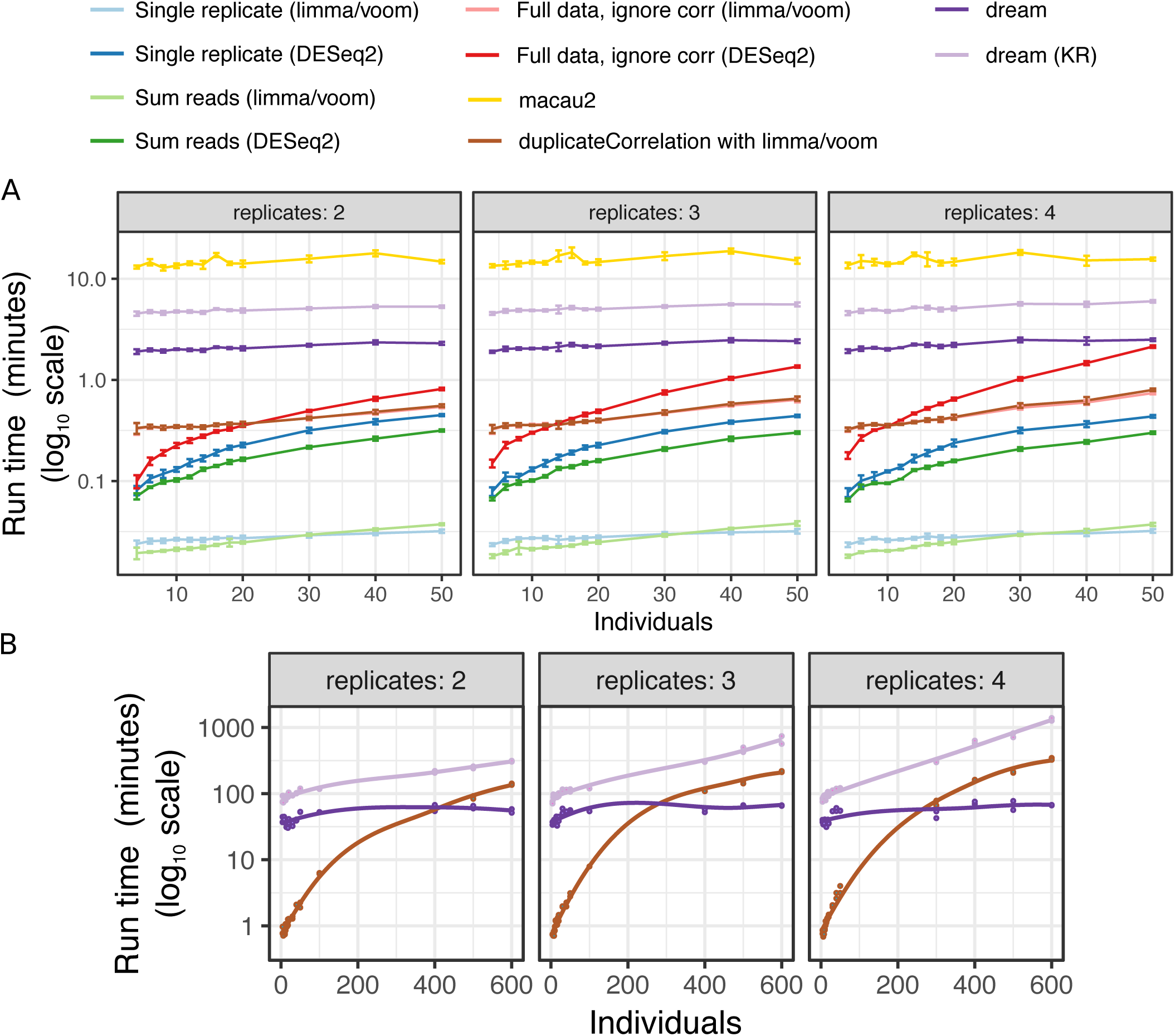
Run time comparison. Run time for was evaluated on the simulated datasets on a 12 core Intel(R) Xeon(R) CPU E5-2620 v3 @ 2.40GHz. **A**) Run time in minutes for each method in the simulation study presented in the previous figures. Averages and standard errors are shown. Analysis was performed using 5 cores. **B**) Run time in minutes for 3 methods on larger simulated datasets using 12 cores. Each combination of individuals, replicates, methods and threads was evaluated on 2 simulated datasets. Lines show loess smoothing. The formula used was: `∼ Disease + (1|Individual)`.

**Figure S 6.**
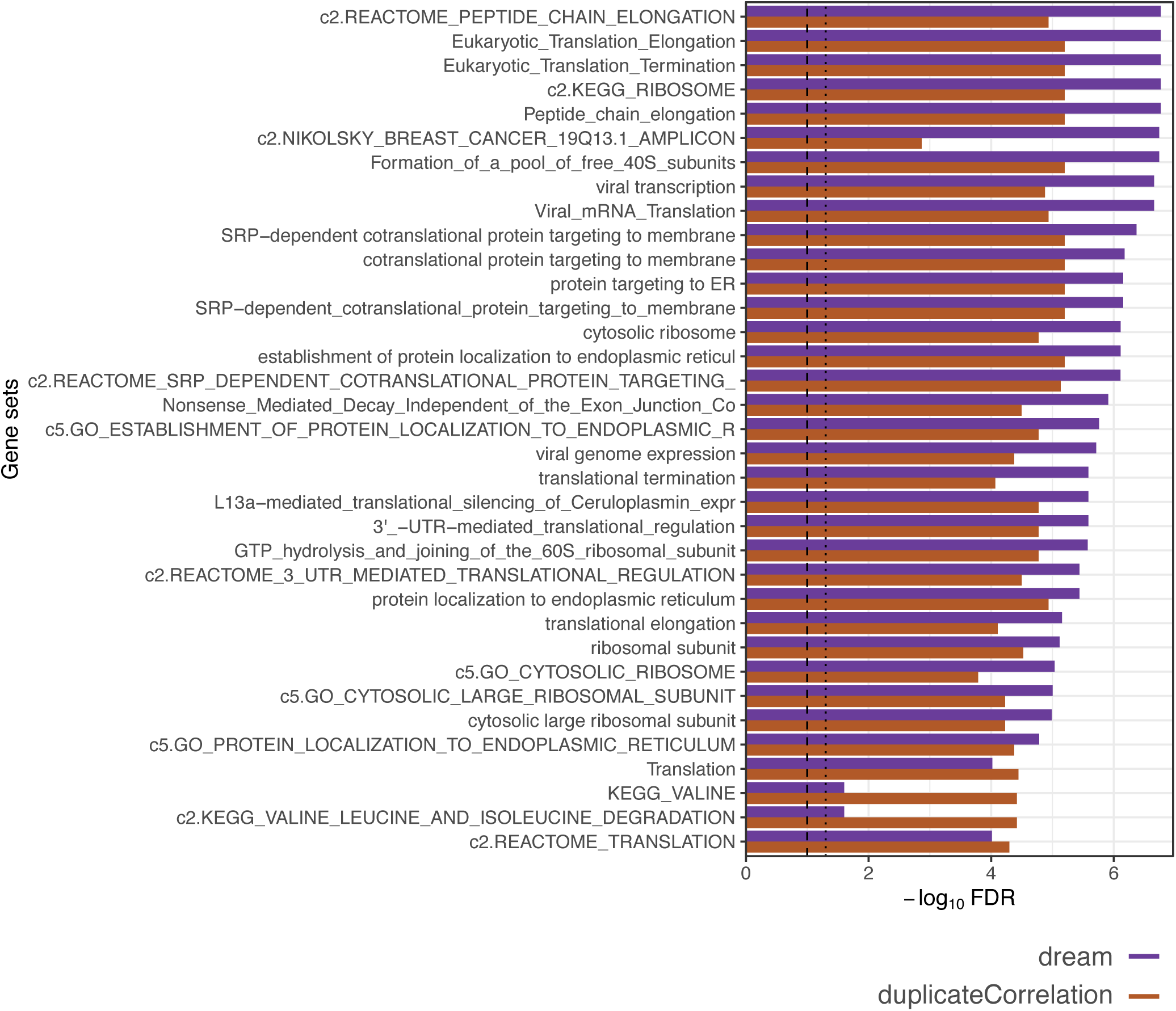
Gene set enrichment FDR for top 30 genesets from differential expression analysis of Braak stage. Enrichment FDRs were computed using t-statistics from dream and duplicateCorrelation.

**Figure S7.**
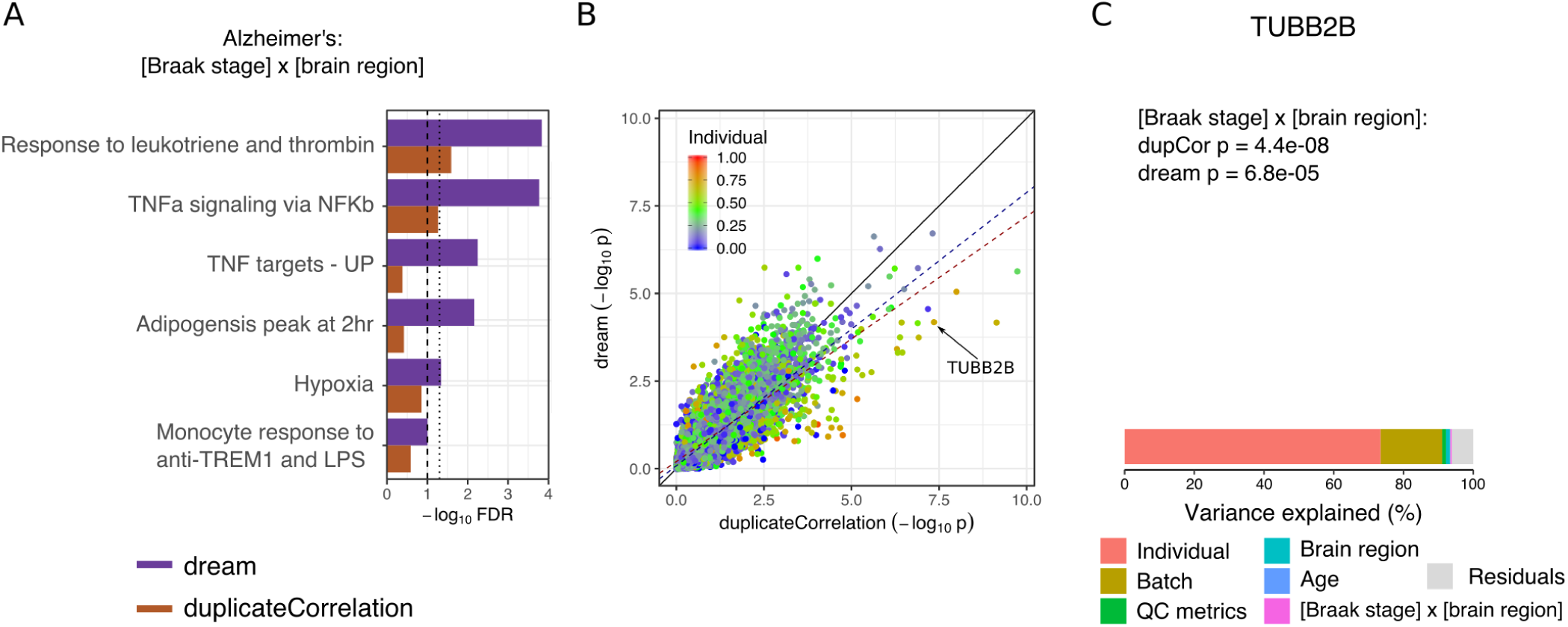
Application to transcriptome data from Alzheimer’s disease. **A**) Gene set enrichment FDR for genes associated with test of Braak stage by brain region term with 4 coefficients. Results are shown for dream and duplicateCorrelation. Lines with broad and narrow dashes indicate 10% and 5% FDR cutoff, respectively. **B**) Comparison of −log_10_ p-values from applying dream and duplicateCorrelation. Each point is a gene, and is colored by the fraction of expression variation explained by variance across individuals. Black solid line indicates a slope of 1. Dashed line indicates the best fit line for the 20% of genes with the highest (red) and lowest (blue) expression variation explained by variance across individuals. **C**) Results for TUBB2B. Box plot is omitted because it is identical to Figure 3C. Bar plot of variance decomposition for TUBB2B shows that 73.4% of variance is explained by expression variance across individuals. Since this value is much larger than the genome-wide mean, duplicateCorrelation under-corrects for the repeated measures.

**Figure S8.**
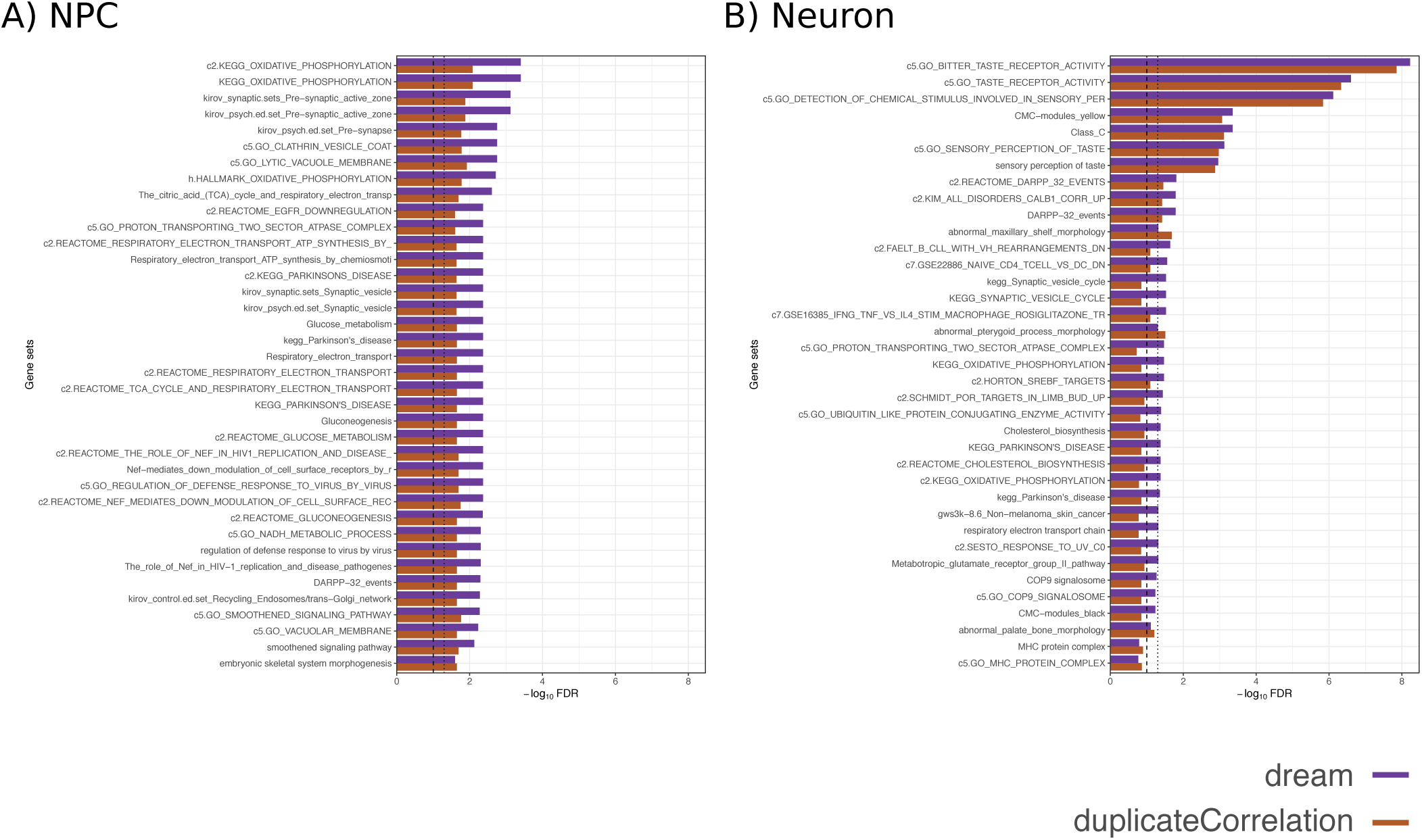
Gene set enrichment FDR for top 30 genesets from differential expression analysis of childhood onset schizophrenia. Enrichment FDRs were computed using t-statistics from dream and duplicateCorrelation analysis of iPSC-derived **A**) neural progenitor cells (NPCs) and **B**) neurons.

**Figure S 9.**
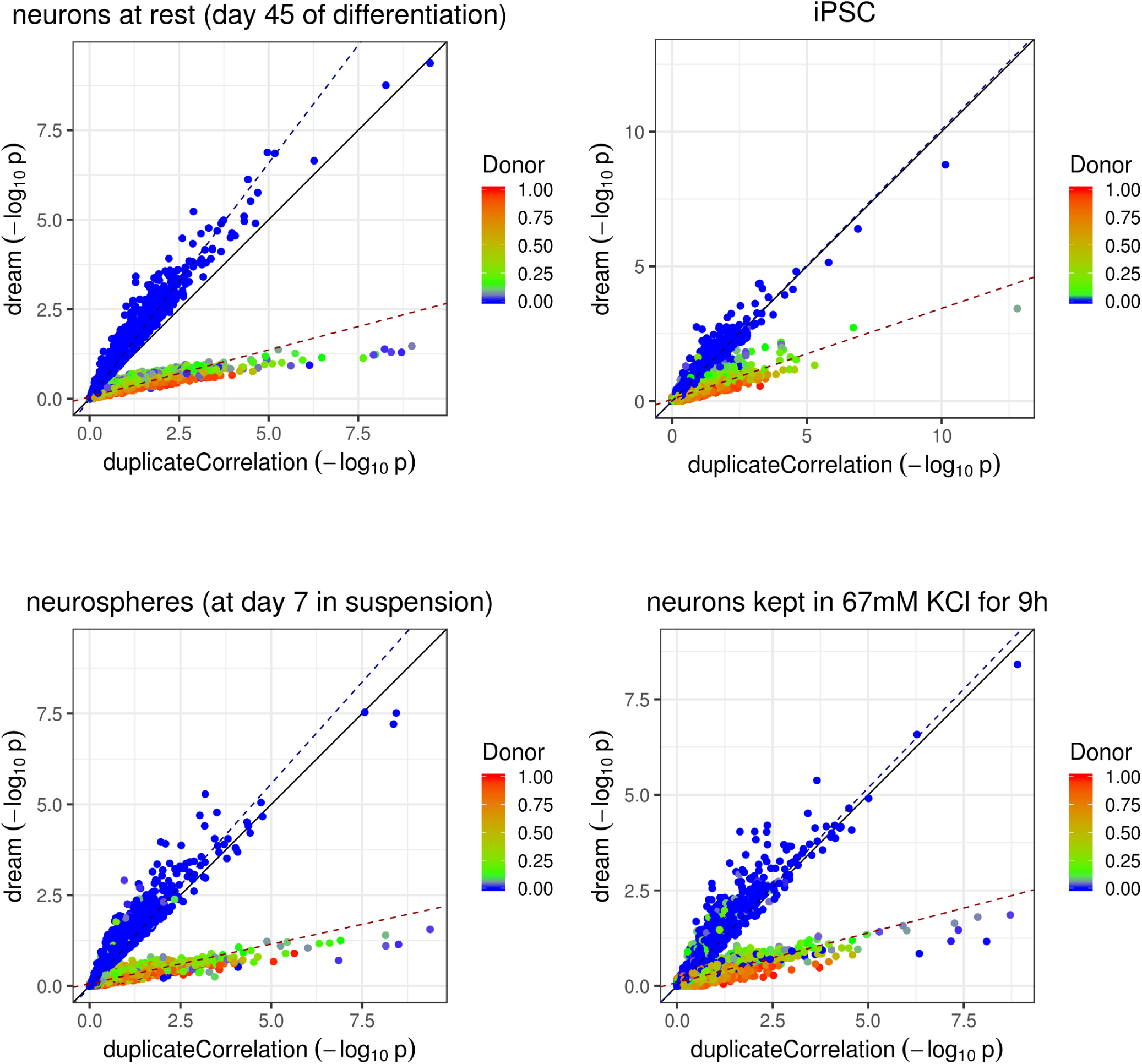
Differential expression analysis of Timothy syndrome compared to controls in four cell types or conditions. Comparison of −log_10_ p-values from applying dream and duplicateCorrelation analyze case/control differences. Each point is a gene, and is colored by the fraction of expression variation explained by variance across individuals. Black solid line indicates a slope of 1. Dashed line indicates the best fit line for the 20% of genes with the highest (red) and lowest (blue) expression variation explained by variance across individuals.

**Figure S 10.**
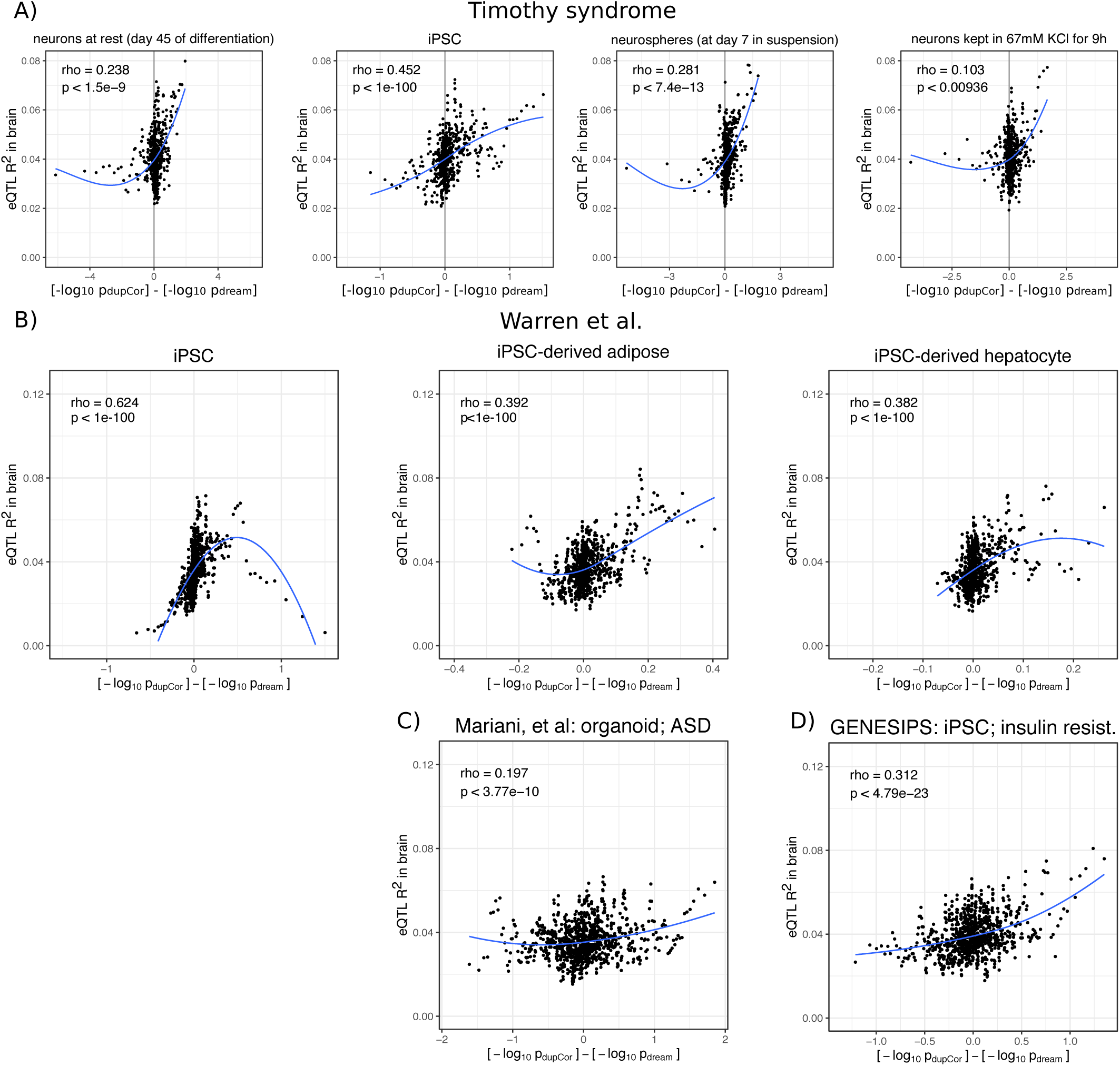
Relationship between differential expression results and genetic regulation. For each gene the fraction of expression variation explainable by cis-eQTLs is compared to the difference in −log_10_ p-value from duplicateCorrelation and dream differential expression analysis. Due to the large number of genes, a sliding window analysis of 100 genes with an overlap of 20 was used to summarize the results. For each window, the average fraction of expression variation explainable by cis-eQTLs (i.e. eQTL R^2^) in brains from the CommonMind Consortium (57) and average difference in −log_10_ p-values from the two methods are reported when differential expression analysis is performed on **A**) Timothy Syndrome in 4 cell types from Pasca, et al. (20) **B**) the SNP rs12740374 in 3 cell types from Warren, et al. (17), **C**) Autism Spectrum in organoids from Mariani, et al. (19), and **D**) insulin resistance in iPSC from Carcamo-Orive, et al. (16). Spearman rho correlations and p-values are shown along with loess curve.

